# Do early life “non-cognitive skills” matter? A systematic review and meta-analysis of early life effects on academic achievement, psychosocial, language and cognitive, and health outcomes

**DOI:** 10.1101/115691

**Authors:** Lisa G. Smithers, Alyssa C. P. Sawyer, Catherine R. Chittleborough, Neil Davies, George Davey Smith, John Lynch

**Affiliations:** School of Public Health, University of Adelaide, Australia; Robinson Research Institute, University of Adelaide, Australia; School of Social and Community Medicine, University of Bristol, Barley House, Oakfield Grove, Bristol, BS8 2BN, United Kingdom; Medical Research Council Integrative Epidemiology Unit, University of Bristol, BS8 2BN, United Kingdom

**Author notes:** Equal first authors (Lisa G. Smithers, Alyssa C. P. Sawyer).

## Abstract

**Background:** Success in school and the labour market is due to more than just high intelligence. Associations between traits such as attention, self-regulation, and perseverance in childhood, and later outcomes have been investigated by psychologists, economists, and epidemiologists. Such traits have been loosely referred to as “non-cognitive” skills. There has been no attempt to systematically assess the relative importance of non-cognitive skills in early life on later outcomes.

**Methods:** The systematic review protocol was registered with the International Prospective Register for Systematic Reviews (PROSPERO, CRD42013006566) in December 2013. We systematically reviewed electronic databases covering psychology, education, health and economics for articles published from database conception until September 2015. Titles and abstracts were screened for eligibility, and from eligible articles data was extracted on study design, sample type and size, age of participants at exposure and outcome, loss to follow up, measurement of exposure and outcome, type of intervention and comparison group, confounding adjustment and results. Where possible we extracted a standardised effect size. We reviewed all studies and rated their evidence quality as ‘better, weak, or poor’ on the basis of study design and potential for confounding, selection and measurement bias.

**Results:** We reviewed 375 studies and provided interpretation of results from 142 (38%) better quality studies comprising randomised controlled trials, quasi-experimental, fixed effects including twin studies, longitudinal and some cross-sectional designs that made reasonable attempts to control confounding. In the academic achievement category outcomes were reported in 78 publications of better quality studies which were consistent with 0.1-0.2 SD effects.

Psychosocial outcomes were reported in 65 better quality studies consistent with effects of 0.3-0.4 SD. For the language and cognitive category there were 42 publications reporting better quality studies consistent with effects of 0.3-0.4 SD. For physical health, results across only eight better quality studies were inconsistent but centred around zero. Analysis of funnel plots consistently showed asymmetric distributions, raising the potential of small study bias which may inflate these observed effects.

**Conclusions:** The evidence under-pinning the importance of non-cognitive skills for life success is diverse and inconsistent. Nevertheless, there is tentative evidence from published studies that non-cognitive skills associate with modest improvements in academic achievement, psychosocial, and language and cognitive outcomes with effects in the range of 0.2-0.4 SD. The quality of evidence under-pinning this field is generally low with more than a third of studies making little or no attempt to control even the most basic confounding (endogeneity) bias. The evidence could be improved by adequately powering studies, and using procedures and tools that improve the conduct and reporting of RCTs and observational studies. Interventions designed to develop children’s non-cognitive skills could potentially improve opportunities, particularly for disadvantaged children. The inter-disciplinary researchers interested in these skills should take a more rigorous approach to determine which interventions are most effective.

## INTRODUCTION

It is forty years since economists Bowles and Gintis^1^, in their critique of the US education system, pointed to the importance of skills for labour market success beyond those captured by intelligence, abstract reasoning and academic achievement in literacy and numeracy. They used the term “non-cognitive personality traits” (pg., 116) and pointed to motivation, orientation to authority, internalization of work norms, discipline, temperament, and perseverance as characteristics that influenced life success. While it may be intuitive that there is more to success in life than high intelligence, there has been no attempt to systematically assess the research evidence on the importance of different types of non-cognitive skills. We recognize there is no neat conceptual dichotomy separating cognitive from some non-cognitive skills, but for the purposes of this review we collectively label the diverse set of factors represented in the literature as “non-cognitive” skills. This literature includes studies that either manipulated non-cognitive skills through randomised controlled trials (RCTs) and quasi-experimental designs, or observed non-cognitive skills through longitudinal or cross-sectional studies. These non-cognitive skills include attention, executive function, inhibitory control, self-control, self-regulation, effortful control, emotion regulation, delay of gratification, and temperament (see Glossary in Supplementary Material for definitions of these constructs). We sought to provide the first systematic representation of research into non-cognitive abilities and behaviours. The need for such a systematic review is driven by the fact that these abilities are being considered by policy makers to underpin early life interventions^2^, beyond cognitive abilities (intelligence or IQ) and academic achievement (literacy, numeracy).

### The policy motivation to improve early life non-cognitive skills

This body of research spans disciplines including psychology, sociology, economics, health and education. It is also of great policy interest to governments in many countries,^2,3^ who wish to sustain future economic productivity and social inclusion, by investing significant resources to bolster the development of human capabilities in early life,^4^ especially for disadvantaged children. The investment logic is that children who develop cognitive and non-cognitive skills early in life will have better outcomes later in life. These outcomes include school readiness, educational attainment, labour market success, better mental and physical health, ultimately leading to a more skilled, healthy and productive workforce. Initial investments beget skills that enable future skills given sustained investments. Non-cognitive skills, such as being able to sustain attention, may be especially important in this regard because they can scaffold later development of cognitive and non-cognitive abilities. If these skills are not developed early in life, then it can be extremely difficult and expensive to compensate later in life, and this reduces returns on later investments.^5^

### Diversity of non-cognitive skills

Since 2000, there has been a 400% increase in publications using keywords describing a variety of non-cognitive skills (Supplementary Figure S1). Several constructs comprise the set of non-cognitive skills reflected in this literature, including externalising behaviours and academic motivation,^6^ responsibility and persistence,^7^ temperament, sociability and behaviour problems,^8^ locus of control and self-esteem,^9^ and attention and socio-emotional skills.^10^ Executive functions^11^ or cognitive control skills (e.g., aspects of how children deploy their cognitive abilities through inhibitory control and attention) may be closely related to cognitive skills, but are also distinguished from IQ, literacy and numeracy.^12^ Personality traits such as self-esteem, patterns of thoughts, feelings, and behaviours that include perseverance, motivation, self-control, and conscientiousness have also been considered as non-cognitive or quasi-cognitive characteristics.^13^ The term “character skills”^14^ has been used to promote the potential malleability of non-cognitive skills in contrast to the notion of personality traits that are thought to be more stable. Heckman and Kautz label these as “soft skills”.^6^ Despite the conceptual complexity and potential overlap of some constructs, many different non-cognitive or personality or character or soft skills are represented in the literature. They have been the target of interventions, especially in early life when these traits are thought to be especially malleable,^15^ and for disadvantaged children, who may benefit most.^5^ Interventions to improve non-cognitive skills may directly improve outcomes,^6,14^ or indirectly, through cognitive ability or other mechanisms. For instance, our own longitudinal analyses in three large population-based cohorts in the UK and Australia showed cognitive abilities were more important than non-cognitive skills in explaining socioeconomic inequalities in academic achievement early in life and that non-cognitive skills were only weakly associated with cognitive ability.^16^

### Evidence for later importance of early life non-cognitive skills

Non-cognitive abilities have been associated with a number of outcomes in later life including mental health^17,18,^ physical health^19^, school readiness and academic achievement^20,21,^ crime^22^, employment and income^9^, and mortality.^23^ Evidence from RCTs suggests that preschool interventions that improve school readiness may do so in part by increasing children’s ability to self-regulate their attention, emotion and behaviour.^24^ Heckman has argued that interventions to develop these skills, especially in disadvantaged young children have the potential for high rates of return due to their positive effects in multiple life domains.^5^

It is widely accepted that children’s cognitive ability (i.e., intelligence or IQ) associates with academic achievement and later success in adulthood.^25-28^ However, the HighScope Perry Preschool Program started in 1962 suggests other mechanisms may be involved.^29,30^ The intervention provided an active learning program based on Piagetian principles, for disadvantaged 3.5 years old African American children who had IQ scores on the Stanford Binet Test < 85^31^. In analysing the long term outcomes of the trial Heckman et al.,^30^ reported that while initially the intervention increased IQ these increases were not maintained to age 7-8 years. Despite this, children who received the intervention went on to enjoy more successful lives in adulthood including greater labour market success, reduced crime involvement and better health.^29,32,33^ While we can find no evidence that the Perry Preschool Program deliberately set out to influence non-cognitive abilities, Heckman and colleagues argued that the intervention resulted in better outcomes for the participants not as a result of increasing their intelligence, but through fostering non-intelligence based socio-emotional “personality” skills^30^ ^(p. 2503)^. This is not dissimilar to the observations of Bowles and Gintis^1^ forty years ago. Schooling does not make children more intelligent, rather, it socializes them into, and rewards, certain characteristics and behaviours that are valued in the labour market.

The aim of this review was to systematically assess all published evidence concerning effects of non-cognitive skills among children up to age eight on later outcomes. We focus on children up to age eight for two reasons. First, these ages are of policy relevance because they are often the focus for early interventions in the health, childcare, preschool, primary school, and child protection sectors. Second, there is evidence that many self-regulatory abilities stabilize around this age^34^. We do not review intervention studies that did not specifically aim to improve non-cognitive skills. Thus, some interventions such as the Perry Preschool^29^ and Abecedarian^35^ programs are not formally reviewed here because we could find no documented evidence that these programs specifically set out to improve non-cognitive abilities, and so were not eligible.

We screened eligible studies and report results on associations between non-cognitive skills under age 8 and four outcome domains - academic achievement, language and cognitive development, physical health, and psychosocial well-being (externalising, internalising behaviour, social skills). In this manuscript we only report results from those studies we judged to be “better” evidence derived from RCTs, quasi-experimental studies, and observational studies that made reasonable attempts to control for confounding (endogeneity) bias. However, all eligible studies were fully reviewed and for completeness are presented in Supplementary Tables S7 and S8.

## METHODS

The systematic review protocol was preregistered with the International Prospective Register for Systematic Reviews (PROSPERO, CRD42013006566) in December 2013 and is available at: http://www.crd.york.ac.uk/PROSPERO/.

### Inclusion criteria

Studies were eligible if they involved non-cognitive abilities of children aged eight years or younger, including executive function (working memory, cognitive flexibility, inhibitory control and attention), effortful control, emotional regulation (emotional reactivity), persistence, conscientiousness, attention, self-control, impulsivity and delay of gratification. Interventions that had general developmental goals were included if they specifically stated an aim related to improving any non-cognitive abilities. Only studies reporting original research were included. Studies involving non-cognitive characteristics in clinical subgroups (e.g., those already diagnosed with problems such as attention-deficit/hyperactivity disorder) were excluded because we were interested in effects of non-cognitive characteristics among developmentally normal healthy children.

We grouped outcomes into four categories - academic achievement (including literacy, numeracy and school readiness), language and cognitive development (including intelligence and language), psychosocial well-being (including mental health problems such as internalising and externalising problems including hyperactivity, social competence, and classroom behaviour), and health (including obesity and injury).

### Literature Search

We searched four electronic databases for articles published from database conception until September 2015: Pubmed, PsycINFO, Embase, and Business Source Complete. These databases were chosen because of their broad coverage of psychological, education, health and economic literature. The search strategy for each database is included in Supplementary Table S1. Search terms were tailored to each database and pilot tested. Study outcomes were not included as search terms to capture all published outcomes associated with non-cognitive abilities. Searches were not restricted by language. Authors of non-English articles were contacted for details or translations. Authors of conference abstracts, editorials and theses were contacted to obtain full text articles. Hand searching of relevant reviews^15,36-38^, our own libraries, and references cited in all RCTs and quasi-experimental interventions was conducted to identify further studies.

### Screening

The titles and abstracts of all articles were screened for eligibility (by AS, LS, CC and TN). To ensure consistency of searching, the first 300 references were searched as a group by all authors and subsequent references were searched independently (Kappa values for agreement were >0.80). Where eligibility was not able to be determined by the title or abstract the full text was reviewed, and when eligibility was unclear this was resolved by group consensus.

### Data extraction

The following information was systematically extracted from each article using a standardized form created by the authors. It included: study design, population-based or convenience sample, age of participants at exposure and outcome measurement, sample size and loss to follow up, measurement of exposure and outcome, type of intervention and comparison group, confounding adjustment and results. To be categorised as a population-based study the publication needed to report some intent and procedure to sample from a defined population base. Where studies did not report age but did report school grade, ages were approximated on the basis of knowledge of school attendance age in the country of interest. LS, JL, CC, AS, TG and TN extracted data from articles. ND independently (i.e., blinded to assessments of other authors) reviewed the data extraction for 15% of all studies, including all intervention studies, and consensus was reached for the very small number of discrepancies.

Where possible we extracted a standardised ‘beta’ coefficient or standardised effect size to have a unit free way of comparing effects across exposures and outcomes (i.e., the difference in SD units between intervention and control groups, or the effect of a 1 SD increase in exposure on an outcome in observational studies). When unstandardized coefficients were reported, where possible we calculated standardised effect sizes to allow comparability of effects across the studies. When a standardized effect size could not be calculated (i.e., SDs for exposure and outcome were not reported) we reported the unstandardized effect sizes.

### Screening to assess risk of bias

The authors JL, LS and AS reviewed all studies and rated their evidence quality as ‘better, weak, or poor’ on the basis of study design and confounding adjustment (Table 1). For RCTs, the risk of bias was assessed using the Cochrane Collaboration Risk of Bias Assessment Tool^39^ (Supplementary Table S6). Observational studies using fixed-effects regression (i.e., twins, and within-individual change) or adjustment for strong common causes of the exposure – outcome association (including proxies for these such as baseline measures of the outcome, or child’s cognitive ability) were rated as better evidence. Studies that inadequately adjusted for confounding, or used subsamples of populations (e.g., selecting participants on the basis of risk for the outcome) were assigned to the ‘weak’ evidence category. Studies that made little or no attempt to adjust for confounding were assigned to the ‘poor’ evidence category. Here we only report results from studies that met the definition of ‘Better evidence’. However, all weak and poor evidence studies were reviewed and appear in Supplementary Tables S7 and S8.

### Data synthesis

We synthesised the information on effect sizes (when presented in the original publication or calculated by us) by undertaking random effects meta-analysis using inverse variance weighting. When no measure of variance was reported we calculated confidence intervals from p values.^40^ It was common for studies to not report variance or exact p values. To overcome this problem for conducting meta-analyses using inverse variance weighting we were forced to make assumptions about p values to calculate confidence intervals. If exact p values were not reported we used the following classification: if p was reported as less than a specific value we assumed p equalled that value, e.g. if p was reported as p<0.01 we assumed p=0.01 for the purpose of calculating confidence intervals; if p was reported as greater than a specific value, we assumed that p=that value + 0.05. We summarised the meta-analyses in a series of forest plots that included effect sizes for individual publications according to study design (cluster, individual, quasi-experimental, longitudinal, cross-sectional) and outcome (Supplementary Figures S2-S18). To reduce bias that may have arisen from studies reporting multiple measures of the same outcome, we obtained an overall estimate across all of the reported measures. For example, if a study reported three different measures of academic achievement we meta-analysed those three estimates to get an overall effect. A final estimate for each outcome was obtained by meta-analysing these summary effects (Figure 2). Additionally, we examined asymmetry of the published evidence by generating funnel plots of effect size against inverse of study size (Supplementary Figures S19-S22) and calculated the summary Egger regression coefficient and P value indicating the degree of asymmetry. To include information on length of follow up we graphed each publication according to length of follow up, effect size and study size (Supplementary Figures S23-S28). The size of the icon in Supplementary Figures S23-S28 corresponds with small (n<100), medium (n 100-500) and large (n>500) studies. The length of the line displays the duration of follow-up.

## RESULTS

The systematic search identified 9618 articles from electronic and hand-searched sources. After removing duplicates and assessing eligibility, 375 articles were included and presented in a PRISMA^41^ flowchart (Figure 1). There were 41 (11%) publications involving RCTs and non-randomised quasi-experimental interventions (Table 1). Below we report this group of studies as Experimental/Quasi-experimental studies (EQIs). Observational studies (including twins) accounted for 89% of all publications examining non-cognitive abilities. Of these outcomes, 46% reported on psychosocial and 32% academic achievement outcomes. Individual publications may have reported multiple outcomes across the domains.

Table 1 shows that of the 375 eligible studies, only 38% (n=142) were rated as “better” evidence, 20% classified as weak and 42% as poor, where there was effectively no attempt to control confounding. The Better evidence category does not imply that all of the studies in this category would be considered “strong” evidence in terms of their design and analysis. For example, some of the EQIs included in Better evidence did not receive high quality ratings according to the Risk of Bias tool (Supplementary Table S6). We extracted and reported results from the 142 Better quality evidence publications (Supplementary Tables S2-S5). This information is summarised in Figure 2 and Supplementary Figures S2-S18, S23-S28, which display all studies where an effect size could be calculated.

### Academic Achievement Outcomes

Academic achievement outcomes mostly comprised reading, writing and numeracy, and were most commonly measured by the Woodcock Johnson psycho-educational battery. Figure 2 shows the effect sizes ranged from 0.16 (0.10-0.23) for academic achievement and school readiness to 0.08 (0.06-0.10) for literacy. They are presented in detail in Supplementary Table S2. They are summarised in forest plots in Supplementary Figures S2-S4, and in graphs in Supplementary Figures S23-S24.

#### EQIs

There were 22 publications reporting ten cluster (school or class) RCTs, nine individual RCTs, one where the unit of randomisation was unclear, and two quasi-experimental intervention studies. These EQIs involved interventions delivered in usual preschool classes, special classes and groups additional to usual curriculum, at home, or a combination of these. Interventions ranged from training specific abilities (e.g. executive functions) to interventions that included several components. The interventions included teacher-delivered curriculum, teacher training to improve classroom behavioural management, and training parents in game-based activities. There was about twice as many EQI publications concerning teacher-delivered curricula than EQIs including both parent and teacher components. Average age at the time of intervention was ~4.3 y. The median follow up time was under 1 year. Reasonably sized cluster RCTs (i.e. average total n>200) rarely reported effect sizes greater than 0.2-0.3 SD. The individually randomised trials were small (n≈30) and tended to have larger effect sizes up to ~1 SD. However, these effects were not consistent across outcomes or maintained with longer follow up. Figures S2 and S3 show the evidence from EQIs was consistent with beneficial effects to literacy and numeracy in the order of 0.1 SD (cluster RCT), 0.5 SD (individual RCT), and 0.6 SD (quasi-experimental).

#### Observational

There were 48 longitudinal (including four fixed effects analysis) and eight cross-sectional publications, with three publications reporting results from multiple cohort studies. On average, non-cognitive abilities were measured at age 4.7 y with the average follow up of longitudinal studies occurring 1.7 years later and longest follow up was 10.2 years. Study sizes ranged from 41 to 21,260. The measures of non-cognitive abilities included attention, executive function, inhibitory control, self-control, self-regulation and effortful control assessed via teacher-report, parent report and objective tests such as the Continuous Performance Task, Head Toes Knees Shoulders task and Stroop-like tasks. Effect sizes were heterogeneous, ranging from null to ~0.8 SD. There was little evidence to conclude that any one measurement tool, measurement method (objective or subjective) or underlying non-cognitive construct was consistently associated with academic achievement. Figures S2 and S3 show effects from observational studies consistent with ~0.1 SD.

### Psychosocial Outcomes

Psychosocial outcomes included mental health problems (internalising and externalising behaviour), social skills, and aspects of school readiness, such as learning engagement. Figure 2 shows the effect sizes ranged from 0.14 (0.09-0.18) for social skills to 0.35 (0.06-0.10) for psychosocial aspects of school readiness. They are presented in detail in Supplementary Table S3. They are summarised in forest plots in Supplementary Figures S5-S9, and in graphs in Supplementary Figures S25-S26. Studies were not consistent in scoring of psychosocial outcomes, i.e. higher scores could indicate worse or better functioning. To aid reader’s interpretation of the results, we have converted all effects to be in the same direction so that positive effects indicate better psychosocial outcomes. However, Supplementary Table S3 presents the results as originally reported in individual publications.

#### EQIs

There were 28 publications reporting 14 cluster RCTs in classrooms, and 12 individual RCTs where the intervention was delivered in schools, sports classes, at home, or in community-based settings. Two quasi-experimental intervention studies involved teacher training or comparing four different preschool programs. Content of the interventions was diverse and included teacher-delivered curriculum sometimes specifically targeting self-regulatory abilities, parent-teacher engagement, teacher training to improve classroom behaviour, training parents in game-based activities, parental Motivational Interviewing and behaviour management, and martial arts. Average age at the time of intervention was ~4.3 years with median follow up time 0.8 years. Intervention groups ranged in size from n=16 to 314 for the individually-randomised trials and n=20 to ~3,350 for cluster RCTs (the largest RCT did not report the exact intervention number). For externalising outcomes, the cluster RCTs reported strong, weak and null effects. For social skills outcomes, cluster RCTs and the non-randomised interventions were also inconsistent. Individual RCTs also reported strong, weak, null, and detrimental effects. Across RCTs there was no consistent evidence favouring one mode of intervention delivery over another. The two largest cluster RCTs that had both a teacher and parent engagement component,^42,43^ which theoretically might be expected to generate the largest effects, only found effects on three of the ten outcomes studied. Figures S5 to S9 show the evidence from EQIs was consistent with beneficial effects across the range of psychosocial outcomes in the order of 0.2 to 0.3 SD.

#### Observational

Four reasonably-sized twin studies that combined MZ and DZ twins (n ranged from 242-410 pairs) of children aged ~2-8 reported phenotypic correlations between non-cognitive abilities and internalising problems of 0 to −0.3, and −0.1 to −0.6 for fewer externalising problems. There were 26 publications of longitudinal studies ranging in size from 49 to 7,140, and seven publications of cross-sectional studies ranging from 42 to 971. On average, non-cognitive skills were measured at age 4.4 years and follow up of the longitudinal studies occurred 2.0 years later. Exposures included attention, executive function, inhibitory control, self-regulation, emotion regulation, delay of gratification, effortful control and temperament, and were assessed by teacher-report, parent report and objective tests. Figures S5 to S9 show effects from observational studies consistent with ~0.1 to 0.2 SD.

Studies of psychosocial outcomes were the most heterogeneous in terms of measuring exposures and outcomes, complicating interpretations of overall effect estimates. There was little evidence that attention (four studies), executive function (two studies) and delay of gratification (one study) affected psychosocial outcomes. For inhibitory control, selfregulation, emotional regulation and temperament, there was some evidence of small to moderate effects (0.1 to 0.6 SD) on social skills and mental health problems. For effortful control, evidence was mixed, ranging from null to moderate effects on psychosocial outcomes.

### Language & Cognitive Outcomes

Language and cognitive outcomes were typically assessed by measures of overall intelligence (such as the Wechsler suite of intelligence tests), verbal and performance intelligence, and language development including expressive and receptive vocabulary (such as the Peabody Picture Vocabulary Test). Figure 2 shows the effect sizes ranged from 0.03 (-0.03-0.09) for verbal IQ to 0.33 (-0.10-0.76) for general cognitive development. They are presented in detail in Supplementary Table S4. They are summarised in forest plots in Supplementary Figures S10-S16, and in graphs in Supplementary Figures S27-S28.

#### EQIs

There were 23 publications reporting 19 RCTs (two interventions were reported in six publications) and four quasi-experimental intervention studies. Of the RCTs, four were cluster (school or class) RCTs, one where the unit of randomisation was unclear, and nine individual RCTs, involving programs delivered in schools or classrooms, at home, in a laboratory setting or a combination of classes and home. Three quasi-experimental interventions involved preschool programs and one involved computerised working memory and inhibitory control training. The content of the interventions was diverse in both delivery and specific focus on non-cognitive ability. Interventions ranged from narrow focused computer-based training to broader content and delivery by teachers in schools plus home visiting with parents. Average age at intervention was 4.2 years, with median follow up of 1 year extending to 16 years. Overall, four reasonably-sized, cluster RCTs (i.e. average n>200) suggested small effects (~0.1-0.3 SD) on language. However, one RCT inconsistently reported effects of 0.15 and 0.25 SD on the same language outcome, using the same sample at age five,^44,45^ and an effect of 0.10 at age 6 in a different publication.^46^ Effects from quasi-experimental studies ranged from null to 0.7 SD. The studies with multiple follow-ups showed early gains in intelligence attenuated with age. Seven of the nine individual RCTs suggested small or no effects on language and intelligence. Two small (n=66, 25) convenience sample RCTs showed larger effects on intelligence (0.4 and 0.6 SD respectively). Figures S10 to S16 show the evidence from EQIs was consistent with effects of null to 0.1 SD for IQ and 0.1-0.2 SD for vocabulary.

#### Observational

A twin study combining MZ and DZ twins (n=237 pairs) showed phenotypic correlations between executive function (predominantly attention) and intelligence at age 12 ranging from 0.1-0.4, with stronger associations for teacher and parent report versus objective assessments. A smaller study combining MZ and DZ twins (n=56 pairs), showed mother-reported attention (measured with 1 question) was associated with 0.13 SD increase in IQ. There were 12 longitudinal and 6 cross-sectional publications that ranged in effect size from - 0.38 (a cross-sectional convenience sample n=77 examining attention) to 0.56 SD (a crosssectional convenience-sample n=80 examining executive function). On average, exposure was measured at age 3.9 years. The median duration of follow up for the longitudinal studies was 0.6 years and the longest follow up was 6.6 years. Exposures included attention, executive function, self-regulation, effortful control and temperament assessed via parent- and teacher-report questionnaires such as the Child Behavior Questionnaire, and objective tests such as the Continuous Performance Task and the HTKS task. There were no longitudinal studies of the effects of attention on language and cognitive outcomes. There were five convenience sample studies of executive function, one cross-sectional and four with shortterm follow up ranging from one to 18 months where effects ranged from 0.06 to 0.33 SD. There were too few studies to examine effortful control and temperament. Six of the seven publications on effects of self-regulation used the HTKS task and showed small effects on vocabulary. Figures S10 to S18 show effects from observational studies was consistent with that from EQIs with effects of null to 0.1 SD for IQ and 0.1 to 0.2 SD for vocabulary.

### Health Outcomes

There were two small RCTs, one quasi-experimental intervention, and nine observational studies that ranged in size from 117 to >26,000. The results of health studies are presented in Supplementary Table S5 and in forest plots in Supplementary Figures S17 and S18. The most common outcomes were Body Mass Index (BMI calculated as weight in kilograms/ height in metres squared) and injuries. An RCT suggested that an intervention based on mindfulness with a 3-month follow up showed a 0.56 SD improvement in teacher-rated health and physical development. Another RCT involving preschool and home visits started in the early 1960s showed no effect on teenage parenthood. It is difficult to interpret the effect of the quasi-experimental intervention because of an inadequate description of the control group and the outcome. Of the observational studies, the average age at exposure was 4.9 years, and duration of follow up in longitudinal studies ranged from 1.5 to 35 years. Observational studies showed little evidence for associations with injury, diet and cardiovascular risk factors, with some evidence that better inhibitory control was associated with lower BMI, but the effect was small.

### Assessment of Publication Bias

The funnel plots in Figures S19-S22 depict effect sizes according to the standard error of the effect size for all publications and outcome categories. All these plots had asymmetric distributions with Egger regression coefficients P-values all less than < 0.01except for health outcomes but there were only 6 studies able to be included.

## DISCUSSION

We reviewed 375 studies and provided interpretation of 142 (38%) better quality studies comprising RCTs, quasi-experimental, fixed effects (including twin studies), longitudinal and some cross-sectional designs. Overall, there is evidence supporting a role for non-cognitive skills in better academic achievement, psychosocial, and language and cognitive outcomes, but as shown in Figure 2, effects are likely to be modest and of the order ~0.2 SD. Our conclusions are broadly consistent with a recent meta-analysis of observational studies of over 14,000 children^47^ that showed a mean effect size of 0.27 for inhibitory control on academic achievement. However, this meta-analysis did not exclude poor quality studies, and did not explore potential for small study bias. We urge some caution in interpreting our results from the published literature, given analysis of funnel plots clearly demonstrate asymmetry of effect size and study size that may raise the potential of small study bias^48^ and/or that larger scale studies are unable to deliver as intensive interventions as small studies. This may mean the meta-analysed effects reported here are over-estimates that may include a null effect.

### Main findings

#### Academic achievement outcomes

Intervention studies focussed on children’s non-cognitive skills at around 4 years of age with median follow-up under one year. These studies were generally consistent with 0.2-0.5 SD short-term effects on academic achievement. However, larger, higher quality RCTs showed smaller effects from 0-0.4 SD.^24,49-51^ These larger higher-quality RCTs spanned child-focussed interventions on specific domains of non-cognitive skills (e.g. Tools of the Mind), to more teacher-focussed curricula (e.g. Chicago School Readiness), to more multi-dimensional content interventions that included parent, child and teacher (e.g. PATHS). One higher quality observational publication^10^ examined six different cohorts with longer follow-up of 5.5 years and reported weaker effects (0-0.2 SD) than intervention studies. Overall, there was insufficient evidence upon which to base a conclusion about the relative effectiveness of these different modes and mechanisms of intervention on non-cognitive skills. Even within the same study, effect sizes differed according to which aspects of academic achievement were measured. For example, one study showed an effect on numeracy but not literacy.^50^ Similarly, another RCT showed that effects on literacy depended on the component of literacy that was measured^44,45^ and effects on some outcomes faded after one year.^46^

#### Psychosocial outcomes

For psychosocial outcomes, the evidence from RCTs was dominated by studies of externalising problems, with fewer RCTs on social skills and internalising problems. Average age at the time of intervention was around 4 years with median follow up time under one year. Of the higher quality RCTs examining externalising outcomes, publications reported strong,^52^ weak^53^ and null effects in the largest of the RCTs^42^. These variable effects could be due to differences in the focus of intervention, mode of delivery (parent, teacher or both), or problems with implementation fidelity in larger trials. Similarly inconsistent results were reported for EQIs with social skills outcomes. The heterogeneity of effects is mirrored in the twin, longitudinal and cross-sectional studies. A good example of this is the inconsistent results reported in five publications that all used the same data source.^54-58^ Across these five publications, interpretation of the effects of self-regulatory abilities depended on how the exposure (attention, delayed gratification, and inhibitory control) and outcome (social skills, withdrawal, and aggression) was measured. The different measures of attention had different associations with the same social skills outcome.

Inhibitory control was associated with social skills and aggression, but not social withdrawal, whereas the effects of delayed gratification on social skills depended on whether the outcomes were directly observed or from maternal report.

The psychosocial outcome studies are the most diverse in interventions (ranging from martial arts, to Motivational Interviewing and Tools of the Mind), and exposure and outcome measurement. This diversity reflects different approaches to improving children’s psychosocial outcomes, such as supporting parents or helping teachers to manage classroom behaviour. Each approach points to different conceptualisations of where psychosocial problems arise and for how, where and whom to intervene (e.g. teachers, psychologists, community nurses or social workers).

#### Language and cognitive outcomes

The relatively small number of studies in this outcome domain (n=28) produced a wide range of effects. Three reasonably-sized cluster RCTs provide the best estimate of the effect of non-cognitive skills on language and cognitive outcomes.^24,50^ They found small effects of ~0.1-0.2 SD. The larger effect sizes (>0.3 SD) are from a well-designed regression discontinuity study (0.44SD),^59^ a non-randomised intervention (0.55-0.73 SD)^60^ and a small, low quality randomised trial.^61^ However, all these studies were small (ranging from N = 12 to 64) and reported effects that attenuated over time or were inconsistent at different ages. The observational studies provide little evidence that the effects are likely to be bigger than ~0.1-0.2 SD, with five of seven longitudinal studies showing few differences and cross-sectional studies reporting mixed effects (-0.38 to 0.56 SD). The longitudinal studies were dominated by non-cognitive skills measured using the HTKS and the WJ Picture Vocabulary as the outcome, and despite the popularity of these measurement tools, the results indicate no effects on vocabulary outcomes. Thus, noncognitive abilities appear to have small effects on cognitive and language outcomes (≤0.2 SD).

#### Physical health outcomes

We cannot draw reliable conclusions for the effect of noncognitive abilities on health as there were only eight studies of highly diverse outcomes with results spanning null to small effects.

### Limitations of this review

While the compilation of 375 publications was systematic and replicable, our assessment of the quality of the evidence is based on our judgement of the potential for bias. We *a priori* created criteria for bias based on well-established procedures including quality appraisal tools, evidence hierarchies, directed acyclic graphs and content knowledge about potential sources of confounding and selection bias. While this involves an element of subjective judgement, we are confident that any other reasonable assessment of the quality of evidence would not change the overall conclusions of the review presented here.

It is possible that some relevant articles were not included in this review, even though we undertook an extensive search that included multiple databases, numerous search terms, contacting authors of potentially-relevant papers, and hand searching reference lists of published papers. Studies of systematic review methods have shown that the most difficult to find articles are often smaller, are poorer quality and the results are unlikely to unduly influence the findings in an already large systematic review.^62^

### The value of a systematic review

While there have been reviews of some aspects of non-cognitive skills,^2,3,13,14,47,63^ none have been systematic in covering the entire literature, or included screening for evidence quality. It has long been recognised in health and medical research that non-systematic reviews of research enable the selective use of evidence to support a particular argument.^62^ For evidence consumers, who are often not evidence-quality specialists, competing claims about effects of non-cognitive abilities based on particular studies are hard to reconcile without the safety net of a systematic review. We have paid particular attention in this review to issues of quality of the primary evidence. There is little point in summarising evidence that includes obviously flawed studies that can only distort the overall results and reduce the value of the systematic nature of the review.^64^

This review covers the entire inter-disciplinary research field on the development of non-cognitive skills. The scope of the review should minimise ‘cherry picking’ of results to bolster a particular concept, theory or intervention. This is necessary to advance knowledge given the multidisciplinary nature of this field and is central for informing interventions to boost life chances for disadvantaged children. In health sciences, major advances have been made by coming to agreement and attempting, where possible, to harmonise methods for measuring exposures, outcomes, synthesising and reporting of outcomes. This work includes collaborative efforts such as the EQUATOR network (http://www.equator-network.org/). Such efforts are needed to reduce waste in research,^64,65^ and improve reproducibility of scientific findings.^66,67^

### Implications for future research

#### What are the active ingredients of non-cognitive skills?

Research that has examined non-cognitive skills in childhood has spanned many disciplines and research traditions, leading to a large number of constructs and tasks being investigated that are sometimes similar in their definition and operationalisation.^68,69,70^ In 1927, Kelley labelled this the “jangle” fallacy^71^ ^(pg. 64)^ where constructs are given different names but in fact are virtually identical. This idea has been recently raised in regard to the concept of “grit” which apparently closely overlaps with perseverance.^72^ It is not uncommon for the same objective tasks to be used as measures of different conceptualisations of non-cognitive abilities. For example the Continuous Performance Task (aka “Go/No Go” task) is used in executive functioning research as a measure of sustained attention and inhibitory control, but it is also used as a measure of effortful control.^69^ Similarly, the “Head-Toes-Knees-Shoulders Task” has been used to measure both behavioural self-regulation^73^ and executive functioning^74^. The interventions we reviewed attempted to influence many different facets of non-cognitive skills. Policy makers and researchers ideally need to know what the ‘active ingredients’ are, in order to enhance children’s non-cognitive skills, and ultimately, the relative effectiveness of different interventions. There are no strong scientific reasons to favour a specific skill over another, but nevertheless it remains important to better understand what the active ingredient(s) underlying non-cognitive skills might be, if we want to support their development.

#### Mechanisms of action

Theoretically, we might expect that interventions involving both parents and teachers might have larger effects on children’s outcomes. However, a recent meta-analysis of early childhood education programs found little evidence that those with parenting involvement produced larger effects, unless it involved a high dose of home visits.^75^ Of the academic outcomes reviewed here, over half (55%) involved only preschool teachers. In our review, there is little evidence to decide which mode of delivery is best and we can find no evidence of attempts at purposive testing of which way to intervene (e.g. teacher, student, parent or various combinations). Purposive testing of delivery mode has been usefully deployed in the design of an RCT in regard to the nurse home visiting literature showing better effects using trained nurses compared to para-professionals.^76^ Interventions that trained children in more specific skills such as executive function, generally showed small effects (e.g., Tools of the Mind).^50^ Other studies imply that non-specific interventions seem to generate better generalised outcomes,^30^ which may suggest that more holistic programs including multidimensional content, may better support overall child development and broad-based benefits.

#### Head-to-Head Comparisons of Interventions

Comparative effectiveness research has been widely promoted in health and medical science as an important contribution to knowing which treatments are the most effective^77,78^. The potential for interventions on non-cognitive skills to influence outcomes may be enhanced by similar approaches. We could find almost no evidence of these sorts of purposeful comparative studies in this field. Exceptions were: Barnett et al.,^50^ and Blair and Raver^79^ who examined effects of Tools of the Mind intervention in cluster RCTs and both found small effects of ~0.1 SD for vocabulary. This exception highlights the potential value of these comparisons.

### Evidence Quality

#### Follow-up

There is a paucity of literature with long-term follow-up. Studies typically began at age 4, with median follow-up of one year or less, and with almost no studies with follow-up beyond age 10, there is very little evidence addressing effects on medium to longer-term outcomes. This is no doubt due to funding constraints. However, it is frequently argued that non-cognitive skills development in childhood have major impacts on the long-term adult outcomes^5^. Thus, interventions which have short term effects, but few detectable long term effects are unlikely to be cost-effective. Therefore, longer-term follow-up of RCTs is especially important. Many of the claims in the literature are that early interventions on a specific trait or with a particular intervention have major long-term effects, but there is very little evidence to support this assertion.

#### Bias

The quality of RCTs was not ideal and reporting of some details was poor or even absent. No RCTs had a formal pre-registered protocol and two thirds did not explicitly identify primary outcomes (See Supplementary Table S6 on Risk of Bias Tool^45^). This can allow cherry-picking of significant results within studies rather than focus on a single or small number of pre-stated main outcome(s) that the intervention is theoretically, or empirically (based on previous evidence) meant to most influence. One third of RCTs may have had other potential biases, for example, differential participation in the control and intervention groups, and unclear processes for selection of control participants. Eighty-eight percent of RCTs did not adequately report randomisation procedures, 81% did not report concealment of allocation processes and participant flow, and most failed to address missing data. It was common for cluster RCTs to have too few clusters to achieve balance between intervention and control groups and in some it was unclear whether clustering was adequately dealt with in the analysis. Poor reporting made it difficult to fully assess study quality and we strongly encourage researchers, journal editors and reviewers to use tools such as the CONSORT statement (http://www.consort-statement.org/) for reporting, and for RCTs to be preregistered. These are now mandatory requirements for publishing in most leading health and medical science journals.

We also reviewed recent RCTs to count the number of previous RCTs they cited. For the four most recent RCTs published in 2014-2015, there were 16 previous RCTs of non-cognitive skills on academic achievement available to be cited. The highest number of previous RCTs referenced in any of these 2014-15 publications was four.^79^ It could perhaps be argued that these RCTs intervened on different non-cognitive skills so should not necessarily cite studies of other non-cognitive skills. Nevertheless, attention regulation and self-control were common ingredients of almost all of these interventions (Supplementary Table S2), so the impression is that new studies were not being explicitly justified on the basis of what was already known from existing RCTs.

Our review highlights the well-established phenomenon that larger effects are often found among observational and smaller studies, compared with large randomised controlled trials. Although our assessment included only evidence from higher quality studies with better control of confounding, the larger effects from observational studies are likely due to unmeasured and residual confounding.^80^ Smaller effects observed in larger RCTs may also be due to implementation difficulty in maintaining intervention fidelity (delivering the intervention as intended by its developers) in community settings, and publication bias. This is an important issue for practice and policy as it suggests that effects found in small RCTs may be overestimated or even non-existent. For example, when studies are ‘scaled-up’ the results can be inconsistent or attenuate towards the null, perhaps suggesting that fidelity is harder, perhaps because an expert is no longer delivering the intervention.^48^

Of the 334 observational studies reviewed here, over 60% were judged as ‘weak’ or ‘poor’ quality. Of all observational studies, 47% made little or no attempt to adjust for even basic confounding i.e., common causes of non-cognitive ability and the outcome. Problems of endogeneity and confounding are well known, and likely to result in substantial bias of the association of non-cognitive skills and later outcomes. One regrettable consequence of this is that much of the research effort in this field is not able to shed much light on the question of whether non-cognitive skills matter. This is a waste of time and resources, and promulgates low-quality research. To advance understanding of non-cognitive skills in children and their effects on outcomes later in life, there is little point in amassing more small-scale^81^, biased observational or experimental studies that have higher likelihood of failing to be replicated.^65,66,76^

### Conclusion

So, after all the voluminous research included in this systematic review, do early life noncognitive skills matter? Overall, yes, there is evidence supporting a role for non-cognitive skills in better academic achievement, psychosocial, and language and cognitive outcomes, but in general, effects are likely to be modest and of the order ~0.2 SD as they relate to shortterm outcomes. However, we urge caution in interpreting this overall finding as positive, given the asymmetry shown in funnel plots that may reflect bias induced by a preponderance of small studies. We urgently need more robust evidence about which skills are the active ingredient(s) and which outcomes they affect in the longer-term. These results suggest profitable pathways forward to help improve influences on life success beyond the traditional focus on reading, writing and arithmetic, and IQ. But the research community interested in these diverse aspects of non-cognitive skills needs higher quality studies, and an integrated scientific focus to help answer the policy-relevant questions^82^.

## Acknowledgements

We would like to thank Tamara Nuske and Tom Goodwin for their research assistance in collecting, and initially screening eligibility, and in the preparation of Tables and Figures.

## References

1 Bowles, S. & Gintis, H. Schooling in Capitalist America: Educational Reform and the Contradictions of Economic Life. (Basic Books, 1976).

2 OECD. Skills for Social Progress: The Power of Social and Emotional Skills. (OECD Publishing, 2015).

3 Gutman, L. & Schoon, I. The Impact of Non-cognitive Skills for Young People. (Institute of Education, U.K. Cabinet Office, London, 2013).

4 Allen, G. Early Intervention: The Next Steps. An Independent Report to Her Majesty's Government. (UK Government, London, 2011).

5 Heckman, J. J. Skill formation and the economics of investing in disadvantaged children. Science 312, 1900–1902 (2006).

6 Heckman, J. J. & Kautz, T. Hard evidence on soft skills. Labour Economics 19, 451–464, doi:http://dx.doi.org/10.1016/i.labeco.2012.05.014 (2012).

7 Lindqvist, E. & Vestman, R. The Labor Market Returns to Cognitive and Noncognitive Ability: Evidence from the Swedish Enlistment. American Economic Journal: Applied Economics 3, 101–128, doi:10.1257/app.3.1.101 (2011).

8 Cunha, F., Heckman, J. J. & Schennach, S. M. Estimating the Technology of Cognitive and Noncognitive Skill Formation. Econometrica 78, 883–931, doi:10.3982/ECTA6551 (2010).

9 Heckman, J. J., Stixrud, J. & Urzua, S. The effects of cognitive and noncognitive abilities on labor market outcomes and social behavior. Journal of Labor Economics 24, 411–482 (2006).

10 Duncan, G. J. et al. School readiness and later achievement. Developmental Psychology 43, 1428–1446, doi:10.1037/0012-1649.43.6.1428 (2007).

11 Hendry, A., Jones, E. J. H. & Charman, T. Executive function in the first three years of life: Precursors, predictors and patterns. Developmental Review 42, 1–33, doi:http://dx.doi.org/10.1016/i.dr.2016.06.005 (2016).

12 Diamond, A., Barnett, W. S., Thomas, J. & Munro, S. Preschool Program Improves Cognitive Control. Science 318, 1387–1388, doi:10.1126/science.1151148 (2007).

13 Borghans, L., Duckworth, A. L., Heckman, J. J. & Ter Weel, B. The economics and psychology of personality traits. Journal of Human Resources 43, 972–1059 (2008).

14 Heckman, J. J. & Kautz, T. Fostering and Measuring Skills: Interventions That Improve Character and Cognition. (National Bureau of Economic Research, Cambridge, MA, 2013).

15 Diamond, A. & Lee, K. Interventions shown to aid executive function development in children 4 to 12 years old. Science 333, 959–964 (2011).

16 Pearce, A. et al. Do early life cognitive ability and self-regulation skills explain socioeconomic inequalities in academic achievement? An effect decomposition analysis in UK and Australian cohorts. Social Science & Medicine 165, 108–118, doi:http://dx.doi.org/10.1016/i.socscimed.2016.07.016(2016).

17 Eisenberg, N. et al. Relations among maternal socialization, effortful control, and maladjustment in early childhood. Development and Psychopathology 22, 507–525 (2010).

18 Fergusson, D. M., Boden, J. M. & Horwood, L. Childhood self-control and adult outcomes: Results from a 30-year longitudinal study. Journal of the American Academy of Child & Adolescent Psychiatry 52, 709–717 (2013).

19 Evans, G. W., Fuller-Rowell, T. E. & Doan, S. N. Childhood Cumulative Risk and Obesity: The Mediating Role of Self-Regulatory Ability. Pediatrics 129, e68-e73 (2012).

20 Blair, C. & Razza, R. P. Relating effortful control, executive function, and false belief understanding to emerging math and literacy ability in kindergarten. Child Development 78, 647–663 (2007).

21 Mischel, W., Shoda, Y. & Peake, P. K. The nature of adolescent competencies predicted by preschool delay of gratification. Journal of Personality and Social Psychology 54, 687–696 (1988).

22 Moffitt, T. E. et al. A gradient of childhood self-control predicts health, wealth, and public safety. Proceedings of the National Academy of Sciences of the United States of America 108, 2693–2698 (2011).

23 Kern, M. L. & Friedman, H. S. Do conscientious individuals live longer? A quantitative review. Health Psychology 27, 505 (2008).

24 Raver, C. C. et al. CSRP's Impact on low-income preschoolers' preacademic skills: self-regulation as a mediating mechanism. Child Dev 82, 362–378, doi:10.1111/j.1467- 8624.2010.01561.x (2011).

25 Deary, I. J., Whiteman, M. C., Starr, J. M., Whalley, L. J. & Fox, H. C. The impact of childhood intelligence on later life: following up the Scottish mental surveys of 1932 and 1947. Journal of personality and social psychology 86, 130 (2004).

26 Fergusson, D. M., John Horwood, L. & Ridder, E. M. Show me the child at seven II: childhood intelligence and later outcomes in adolescence and young adulthood. Journal of Child Psychology and Psychiatry 46, 850–858, doi:10.1111/j.1469-7610.2005.01472.x (2005).

27 Kuh, D., Richards, M., Hardy, R., Butterworth, S. & Wadsworth, M. E. Childhood cognitive ability and deaths up until middle age: a post-war birth cohort study. International Journal of Epidemiology 33, 408–413, doi:10.1093/ije/dyh043 (2004).

28 Whalley, L. J. & Deary, I. J. Longitudinal Cohort Study Of Childhood Iq And Survival Up To Age 76. BMJ: British Medical Journal 322, 819–822, doi:10.2307/25466668 (2001).

29 Schweinhart, L. J. et al. Lifetime effects: the High/Scope Perry Preschool study through age 40. (2005).

30 Heckman, J. J., Pinto, R. & Savelyev, P. Understanding the mechanisms through which an early childhood program boosted adult outcomes. American Economic Review 103, 2052–2086 (2013).

31 Weikert, D. P. Comparative study of three preschool curricula. Report No., F244, (Washington, D.C., 1969).

32 Schweinhart, L. J. Significant Benefits: The High/Scope Perry Preschool Study through Age 27. Monographs of the High/Scope Educational Research Foundation, No. Ten. (ERIC, 1993).

33 Heckman, J., Moon, S. H., Pinto, R., Savelyev, P. & Yavitz, A. Analyzing social experiments as implemented: A reexamination of the evidence from the HighScope Perry Preschool Program. Quantitative economics 1, 1–46 (2010).

34 Weintraub, S. et al. I. NIH Toolbox Cognition Battery (CB): introduction and pediatric data. Monogr Soc Res Child Dev 78, 1–15, doi:10.1111/mono.12031 (2013).

35 Campbell, F. & Ramey, C. Effects of early intervention on intellectual and academic achievement: a follow-up study of children from low-income families program title: Carolina Abecedarian Project. Child Development 65, 684 (1994).

36 Blair, C. & Diamond, A. Biological processes in prevention and intervention: The promotion of self-regulation as a means of preventing school failure. Development and psychopathology 20, 899–911 (2008).

37 Blair, C. & Raver, C. C. School Readiness and Self-Regulation: A Developmental Psychobiological Approach. Annual review of psychology 66, 711–731 (2015).

38 Diamond, A. Activities and Programs That Improve Children’s Executive Functions. Current Directions in Psychological Science 21, 335–341 (2012).

39 Higgins, J. P. T. et al. The Cochrane Collaboration’s tool for assessing risk of bias in randomised trials. BMJ 343, doi:10.1136/bmj.d5928 (2011).

40 Altman, D. G. & Bland, J. M. How to obtain the confidence interval from a P value. BMJ: British Medical Journal 343 (2011).

41 Liberati, A. et al. The PRISMA statement for reporting systematic reviews and metaanalyses of studies that evaluate healthcare interventions: explanation and elaboration. BMJ 339 (2009).

42 Webster-Stratton, C., Jamila Reid, M. & Stoolmiller, M. Preventing conduct problems and improving school readiness: evaluation of the incredible years teacher and child training programs in high-risk schools. Journal of child psychology and psychiatry 49, 471–488 (2008).

43 Conduct Problems Prevention Research Group. Initial impact of the Fast Track prevention trial for conduct problems: I. The high-risk sample. Journal of consulting and clinical psychology 67, 631–647 (1999).

44 Nix, R. L., Bierman, K. L., Domitrovich, C. E. & Gill, S. Promoting children's social-emotional skills in preschool can enhance academic and behavioral functioning in kindergarten: Findings from Head Start REDI. Early Education & Development 24, 1000–1019 (2013).

45 Bierman, K. L. et al. Promoting academic and social-emotional school readiness: The Head Start REDI program. Child development 79, 1802–1817 (2008).

46 Bierman, K. L. et al. Effects of Head Start REDI on children's outcomes 1 year later in different kindergarten contexts. Child Development 85, 140–159 (2014).

47 Allan, N. P., Hume, L. E., Allan, D. M., Farrington, A. L. & Lonigan, C. J. Relations between inhibitory control and the development of academic skills in preschool and kindergarten: A meta-analysis. Developmental Psychology 50, 2368–2379, doi:10.1037/a0037493 (2014).

48 Egger, M. & Smith, G. D. Misleading meta-analysis. BMJ 311, 753–754 (1995).

49 Brotman, L. M. et al. Cluster (school) RCT of parentcorps: Impact on kindergarten academic achievement. Pediatrics 131, e1521-e1529 (2013).

50 Barnett, W. S. et al. Educational effects of the Tools of the Mind curriculum: A randomized trial. Early Childhood Research Quarterly 23, 299–313, doi:http://dx.doi.org/10.1016/i.ecresq.2008.03.001(2008).

51 Ialongo, N. S. et al. Proximal impact of two first-grade preventive interventions on the early risk behaviors for later substance abuse, depression, and antisocial behavior. American journal of community psychology 27, 599–641 (1999).

52 Raver, C. C. et al. Targeting children's behavior problems in preschool classrooms: A cluster-randomized controlled trial. Journal of Consulting and Clinical Psychology 77, 302 (2009).

53 Shelleby, E. C. et al. Behavioral control in at-risk toddlers: The influence of the family check-up. Journal of Clinical Child and Adolescent Psychology 41, 288–301 (2012).

54 NICHD Early Child Care Research Network. Do children's attention processes mediate the link between family predictors and school readiness? Developmental Psychology 39, 581–593 (2003).

55 Ramani, G. B., Brownell, C. A. & Campbell, S. B. Positive and negative peer interaction in 3- and 4-year-olds in relation to regulation and dysregulation. The Journal of genetic psychology 171, 218–250, doi:10.1080/00221320903300353 (2010).

56 Runions, K. C. & Keating, D. P. Anger and inhibitory control as moderators of children's hostile attributions and aggression. Journal of Applied Developmental Psychology 31, 370–378 (2010).

57 Mintz, T. M., Hamre, B. K. & Hatfield, B. E. The role of effortful control in mediating the association between maternal sensitivity and children's social and relational competence and problems in first grade. Early Education and Development 22, 360–387 (2011).

58 Booth-Laforce, C. & Oxford, M. L. Trajectories of social withdrawal from grades 1 to 6: prediction from early parenting, attachment, and temperament. Dev Psychol 44, 1298–1313, doi:10.1037/a0012954 (2008).

59 Weiland, C. & Yoshikawa, H. Impacts of a pre kindergarten program on children's mathematics, language, literacy, executive function, and emotional skills. Child Development 84, 2112–2130 (2013).

60 Bradley, R. T., Galvin, P., Atkinson, M. & Tomasino, D. Efficacy of an emotion selfregulation program for promoting development in preschool children. Global Advances In Health and Medicine 1, 36–50 (2012).

61 Ford, R. M., McDougall, S. J. & Evans, D. Parent-delivered compensatory education for children at risk of educational failure: Improving the academic and self-regulatory skills of a Sure Start preschool sample. British journal of psychology (London, England: 1953) 100, 773–797, doi:10.1348/000712609x406762 (2009).

62 Egger, M., Juni, P., Bartlett, C., Holenstein, F. & Sterne, J. How important are comprehensive literature searches and the assessment of trial quality in systematic reviews? Empirical study. Health Technology Assessment 7, 76, doi:10.3310/hta7010 (2003).

63 Diamond, A. Executive functions. Annual Review of Psychology 64, 135–168 (2013).

64 Chalmers, I. et al. How to increase value and reduce waste when research priorities are set. The Lancet 383, 156–165, doi:http://dx.doi.org/10.1016/S0140-6736(13)62229-1.

65 Ioannidis, J. P. et al. Increasing value and reducing waste in research design, conduct, and analysis. Lancet 383, 166–175, doi:10.1016/s0140-6736(13)62227-8 (2014).

66 Open Science Collaboration. Estimating the reproducibility of psychological science. Science 349, doi:10.1126/science.aac4716 (2015).

67 Munafò, M. R. et al. A manifesto for reproducible science. Nature Human Behaviour 1, 0021, doi:10.1038/s41562-016-0021 (2017).

68 Duckworth, A. L. & Kern, M. L. A meta-analysis of the convergent validity of selfcontrol measures. Journal of Research in Personality 45, 259–268, doi:http://dx.doi.org/10.1016/i.irp.2011.02.004 (2011).

69 Zhou, Q., Chen, S. H. & Main, A. Commonalities and differences in the research on children’s effortful control and executive function: A call for an integrated model of self-regulation. Child Development Perspectives 6, 112–121 (2012).

70 Shonkhoff, J. P. & Phillips, D. A. in From Neurons to Neighbourhoods. The Science of Early Childhood Development (eds J. P. Shonkhoff & D. A. Phillips) 93–123 (National Academy Press, 2001).

71 Kelley, T. L. Interpretation of Educational Measurement. (World Books, 1927).

72 Credé, M., Tynan, M. C. & Harms, P. D. Much Ado About Grit: A Meta-Analytic Synthesis of the Grit Literature. Journal of Personality and Social Psychology, No Pagination Specified doi:10 1037/pspp0000102 (9016)

73 Ponitz, C. C., McClelland, M. M., Matthews, J. & Morrison, F. J. A structured observation of behavioral self-regulation and its contribution to kindergarten outcomes. Developmental Psychology 45, 605–619 (2009).

74 Cameron, C. E. et al. Fine motor skills and executive function both contribute to kindergarten achievement. Child Dev 83, 1229–1244, doi:10.1111/j.1467- 8624.2012.01768.x (2012).

75 Grindal, T. et al. The added impact of parenting education in early childhood education programs: A meta-analysis. Children and Youth Services Review 70, 238–249, doi:http://dx.doi.org/10.1016/j.childyouth.2016.09.018 (2016).

76 Olds, D. et al. Effects of home visits by paraprofessionals and by nurses: age 4 followup results of a randomized trial. Pediatrics 114, 1560 - 1568 (2004).

77 Iglehart, J. K. Prioritizing Comparative-Effectiveness Research -- IOM Recommendations. The New England Journal of Medicine, NEJMp0904133 (2009).

78 Fiore, L. D. & Lavori, P. W. Integrating Randomized Comparative Effectiveness Research with Patient Care. New England Journal of Medicine 374, 2152–2158, doi:10.1056/NEJMra1510057 (2016).

79 Blair, C. & Raver, C. C. Closing the achievement gap through modification of neurocognitive and neuroendocrine function: results from a cluster randomized controlled trial of an innovative approach to the education of children in kindergarten. PLoS One 9, e112393, doi:10.1371/journal.pone.0112393 (2014).

80 Fewell, Z., Davey Smith, G. & Sterne, J. A. The impact of residual and unmeasured confounding in epidemiologic studies: a simulation study. Am J Epidemiol 166, 646–655, doi:10.1093/aje/kwm165 (2007).

81 Smaldino, P. E. & McElreath, R. The natural selection of bad science. Royal Society Open Science 3, doi:10.1098/rsos.160384 (2016).

82 Watts, D. J. Should social science be more solution-oriented? Nature Human Behaviour 1, 0015, doi:10.1038/s41562-016-0015 (2017).

